# Two key events associated with a transposable element burst occurred during rice domestication

**DOI:** 10.1101/405290

**Authors:** Jinfeng Chen, Lu Lu, Jazmine Benjamin, Stephanie Diaz, C. Nathan Hancock, Jason E. Stajich, Susan R. Wessler

## Abstract

Transposable elements shape genome evolution through periodic bursts of amplification. In this study we exploited knowledge of the components of the *mPing/Ping/Pong* TE family in four rice strains undergoing *mPing* bursts to track their copy numbers and distribution in a large collection of genomes from the wild progenitor *Oryza rufipogon* and domesticated *Oryza sativa* (rice). We characterized two events that occurred to the autonomous *Ping* element and appear to be critical for *mPing* hyperactivity. First, a point mutation near the end of the element created a *Ping* variant (*Ping16A*) with reduced transposition. The proportion of strains with *Ping16A* has increased during domestication while the original *Ping (Ping16G)* has been dramatically reduced. Second, transposition of *Ping16A* into a *Stowaway* element generated a locus (*Ping16A_Stow*) whose presence correlates with strains that have high *mPing* copies. Finally, demonstration that *Pong* elements have been stably silenced in all strains analyzed indicates that sustained activity of the *mPing/Ping* family during domestication produced the components necessary for the *mPing* burst, not the loss of epigenetic regulation.

## Introduction

Eukaryotic genomes are populated with transposable elements (TEs), many attaining copy numbers of hundreds to thousands of elements by rapid amplification, called a TE burst. For a TE to successfully burst it must be able to increase its copy number without killing its host or being silenced by host surveillance. However, because the vast majority of TE bursts have been inferred after the fact – via computational analysis of whole genome sequence – the stealth features they require for success have remained largely undiscovered.

Revealing these stealth features requires the identification of a TE in the midst of a burst. This was accomplished for the miniature inverted-repeat transposable element (MITE) *mPing* from rice^1,2^. MITEs are non-autonomous Class II (DNA) elements that are the most common TE associated with the non-coding regions of plant genes^3^. To understand how MITEs attain high copy numbers despite a preference for insertion into genic regions, a computational approach was used to identify *mPing*, and its source of transposase, encoded by the related autonomous *Ping* element (Fig. 1a)^1^.

**Figure 1.**
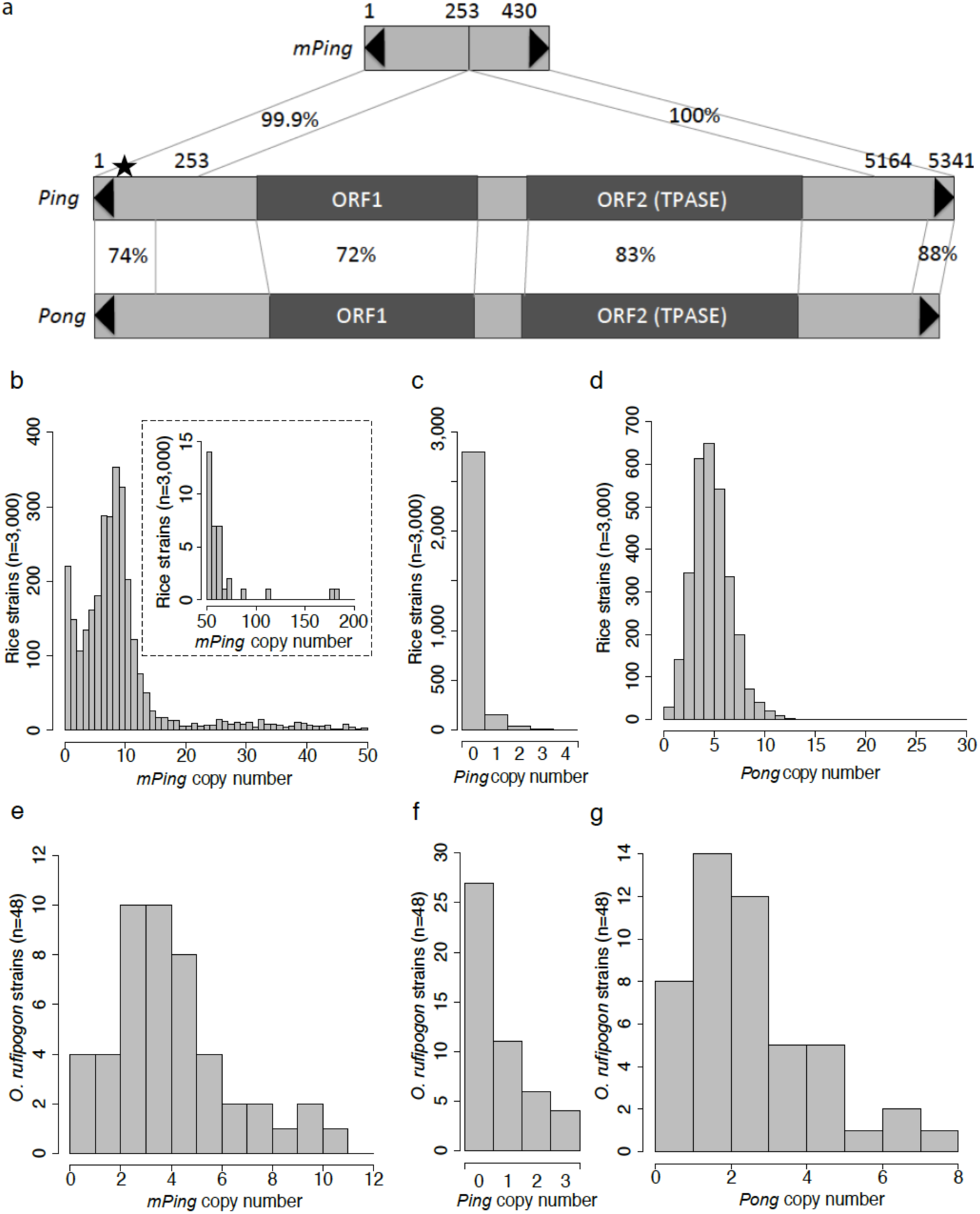
Abundance of *mPing, Ping* and *Pong* elements in 3,000 rice and 48 *O. rufipogon* genomes. **a**, Comparison of structures of *mPing, Ping* and *Pong*. TIRs are indicated by black triangles. Two protein coding genes ORF1 and ORF2 (TPASE) encoded by *Ping* or *Pong* are indicated by dark gray boxes. Homologous regions between elements are connected by lines and percent identities are shown. The black star on *Ping* indicates the +16G/A SNP that differs between *mPing* and *Ping16A*. Copy numbers across the 3,000 rice strains of *mPing* (**b**), *Ping* (**c**), and *Pong* (**d**). The bar plot in the dashed-box in **b** shows strains with more than 50 *mPing* elements. **e**, *mPing* copy number of 48 *O. rufipogon* strains. **f**, *Ping* copy number of 48 *O. rufipogon* strains. **g**, *Pong* copy number of 48 *O. rufipogon* strains.

Ongoing bursts of *mPing* were discovered in four temperate *japonica* strains: EG4, HEG4, A119, and A123, whose genomes were sequenced and insertion sites and epigenetic landscape determined^2,4,5^. These analyses uncovered two features of successful bursts. First, *mPing* targets genic regions but avoids exon sequences, thus minimizing harm to the host^2,5^. Second, because *mPing* does not share coding sequences with *Ping* (Fig. 1a), increases in its copy number and host recognition of its sequences does not silence *Ping* genes, thus allowing the continuous production of the proteins necessary to sustain the burst for decades^4^.

The contributions of two other features to the success of the bursts could not be assessed previously and are the focus of this study. These features are a single SNP at position 16 (+16G/A) that distinguishes *mPing* and *Ping* sequences (Fig. 1a), and a single *Ping* locus (called *Ping16A_Stow*) that is the only *Ping* locus shared by all bursting strains^4^. To understand the origin of these features and their possible role in the burst, we analyzed the presence, sequence, and copy numbers of *Ping* and *mPing* elements in the genomes of 3,000 domesticated rice strains and 48 genomes of their wild progenitor, *O. rufipogon*. Rice has been divided into five major groups or subfamilies that are thought to have originated from distinct populations of the wild progenitor *O. rufipogon* that arose prior to domestication (Table 1)^6,7^. Furthermore, significant gene flow from *japonica* to *indica* and *aus* has been noted previously, reflecting the more ancient origin of *japonica*^6,8^.

**Table 1.** Distribution of *Ping* variants and *Ping16A_Stow*^a^ genotypes in domesticated rice and *O. rufipogon*

| Subgroups | Number of strains | Number of strains with *Ping*<sup>b</sup> | *Ping* variants |  | *Ping16A_Stow* genotypes |  |
| --- | --- | --- | --- | --- | --- | --- |
|  |  |  | Number of strains with *Ping16G* | Number of strains with *Ping16A* | *Stowaway* only | *Stowaway* with *Ping* |
| *O. sativa* | 3,000 | 199 (6.6%) | 31 | 154 | 188 | 11 |
| -*indica* | 1,651 | 20 (1.2%) | 8 | 9<sup>c</sup> | 10 | 0 |
| -*aus/boro* | 189 | 28 (14.8%) | 19 | 0 | 0 | 0 |
| -temperate *japonica* | 250 | 61 (24.4%) | 1 | 61 | 121 | 8 |
| -tropical *japonica* | 335 | 51 (15.2%) | 0 | 51 | 2 | 0 |
| -*aromatic* | 65 | 0 (0%) | 0 | 0 | 0 | 0 |
| -admixed | 510 | 39 (7.6%) | 3 | 33<sup>d</sup> | 55 | 3 |
| *O. rufipogon* | 48 | 21 (43.7%) | 21 | 0 | 4 | 0 |
| -*Or-I* | 13 | 7 (53.8%) | 7 | 0 | 0 | 0 |
| -*Or-II* | 23 | 10 (43.4%) | 10 | 0 | 1 | 0 |
| -*Or-IIIa* | 6 | 2 (33.3%) | 2 | 0 | 3 | 0 |
| -*Or-IIIb* | 6 | 2 (33.3%) | 2 | 0 | 0 | 0 |

<sup>a</sup> *Ping16A\_Stow* is defined as a locus where *Ping* has inserted into the *Stowaway* element on chromosome 1 (2640500-2640502)
<sup>b</sup> “Number of strains with *Ping16G*” plus “Number of strains with *Ping16A*” is less than or equal to “Number of strains with *Ping*” because *Ping* genotypes in some strains cannot be determined from available sequences. An exception is “temperate *japonica*”, where one strain (IRIS\_313-10564) has both *Ping16G* (Chr8:2964281-2964283) and *Ping16A* (Chr6:23521641-23526981).
<sup>c</sup> Eight *indica* strains have *Ping16A* that are located in regions showing evidence of introgression from *japonica* (Seven strains share the locus Chr3:21965880-21965882 and one strain has the Nipponbare *Ping* locus Chr6:23521641-23526981). One *indica* strain has *Ping16A* in a region with *indica* background.
<sup>d</sup> Thirty-one admixed strains have *Ping16A* from *japonica*. Two admixed strains have *Ping16A* that are located in regions with ambiguous origin.

Knowledge of the relationships between the major groups of rice and the populations of *O. rufipogon* have been utilized in this study to better understand the identity and origin of the components necessary for *mPing* bursts. Of particular interest was whether (i) *mPing* bursts could be detected in other strains of wild and/or domesticated rice, (ii) the +16G/A *Ping* SNP and *Ping16A_Stow* originated in wild rice or first appeared after domestication, and (iii) the presence of +16G/A *Ping* SNP and *Ping16A_Stow* correlated with higher *mPing* copy number.

Finally, another potential player that may be implicated in *mPing* bursts, *Pong*, is a focus of this study (Fig. 1a). *Pong* is the closest relative of *Ping* in the rice genome with at least five identical copies in all strains of rice analyzed to date^4,9^. Relevant to this study is that *Pong* encoded proteins catalyzed the transposition of *mPing* in rice cell culture^1^ and in transposition assays in *Arabidopsis thaliana* and yeast^10,11^. However, *Pong* elements do not catalyze *mPing* transposition in planta because all *Pong* copies are effectively silenced and its sequences are associated with heterochromatin^4^. Here we were able to address questions regarding the origin and stability of *Pong* silencing before and during domestication.

## Results

### Detection of *mPing, Ping,* and *Pong* elements

Insertion sites and copy numbers for m*Ping, Ping*, and *Pong* were identified from genome sequences of 3,000 rice strains using RelocaTE2^12^ (see Methods). The paired-end DNA libraries had an average insert size of ∼ 500 bp and were sequenced to a depth of 14-fold genome coverage^13^ which allowed clear distinction between *mPing, Ping, and Pong* elements (Fig. 1a). Sequence analyses identified a total of 27,535 *mPings*, 262 *Pings*, and 12,748 *Pong*s (Fig. 1b-d and Supplementary Table 1). Copy numbers of *mPing, Ping*, and *Pong* elements in each genome were also estimated using a read-depth method (see Methods). Outputs from the RelocaTE2 and read depth methods were well correlated (Pearson’s correlation, *R* = 0.97, *P* < 2.2e-16 for *mPing*; *R* = 0.9, *P* < 2.2e-16 for *Ping*; *R* = 0.66, *P* < 2.2e-16 for *Pong*; Supplementary Fig. 1) suggesting that both methods were robust. Insertion sites and copy numbers for *mPing, Ping* and *Pong* were also identified for 48 *O. rufipogon* strains, but only the read-depth method was used because of the limited insert size of the libraries (Supplementary Table 2). In total, 193 *mPing*s, 23 *Pings*, and 124 *Pongs* were estimated to be present in the 48 *O. rufipogon* strains (Fig. 1e-g and Supplementary Fig. 2).

**Table 2.**
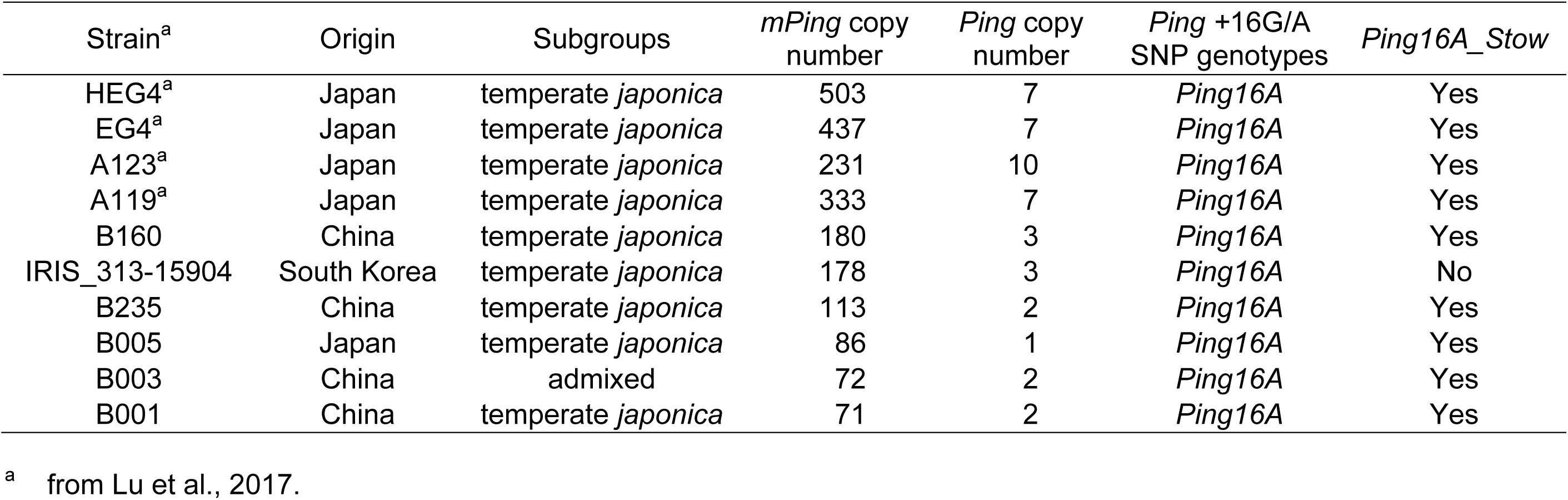
*Ping* copy numbers and genotypes in rice strains with high copy numbers of *mPing*

**Figure 2.**
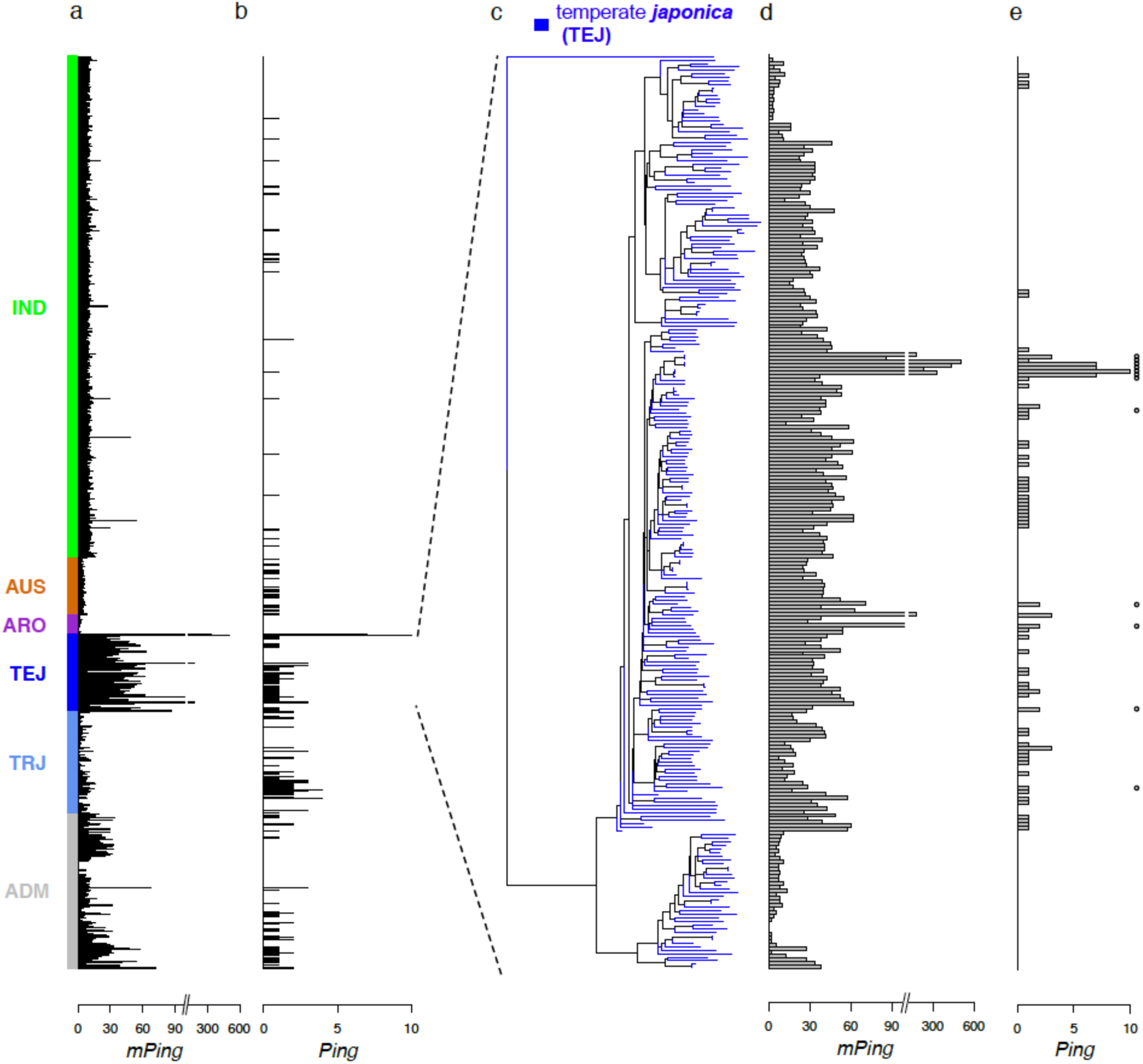
Copy numbers of *mPing, Ping* and *Pong* elements in rice subgroups of the 3,000 sequenced genomes and the four strains undergoing *mPing* bursts (HEG4, EG4, A119, and A123). **a**, *mPing* copy numbers in 3,000 genomes. Colors represent the five major rice subgroups: *indica* (IND), *aus/boro* (AUS), *aromatic* (ARO), temperate *japonica* (TEJ), tropical *japonica* (TRJ), and admixed (ADM). **b**, *Ping* copy numbers in 3,000 genomes. **c**, Neighbor-joining tree of 250 temperate *japonica* strains using genome-wide SNPs. **d**, *mPing* copy number of temperate *japonica* strains. **e**, *Ping* copy number of temperate *japonica* strains. Strains that the *Ping16A_Stow* locus are noted with open circles.

### Copy number variation of *mPing* and *Ping* elements in domesticated and wild rice

None of the 3,000 rice strains analyzed in this study have more *mPing* elements than the 231-503 copies found in the four temperate *japonica* strains (HEG4, EG4, A119, A123) in the midst of *mPing* bursts^4^. Of the 3,000 rice strains, 2,780 (92.7%) contain *mPing*, with an average of about 9 elements per strain (Fig. 1b). Temperate *japonica* strains do, however, have significantly more *mPing* elements (∼30.5/strain) than tropical *japonica* (∼2.6/strain), *indica* (∼8.2/ strain), or *aus/boro* (∼3.8/strain) (Supplementary Table 3 and Supplementary Fig. 3). All *O. rufipogon* strains have *mPing* elements with copy numbers ranging from 1-11 (Fig. 1g and Supplementary Fig. 2).

**Figure 3.**
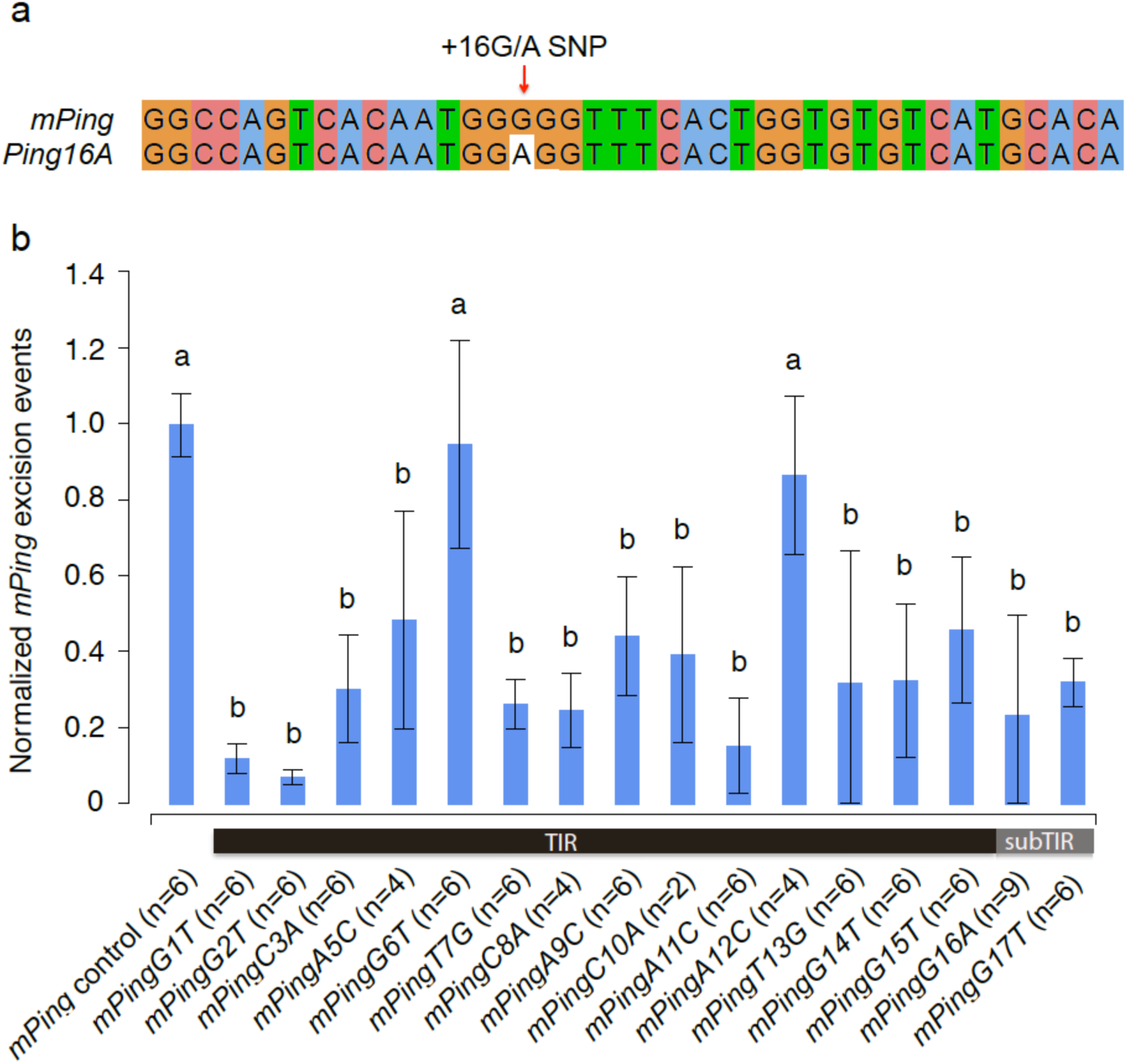
Transposition frequency of *mPing* variants in the yeast assay. **a**, Sequence alignment of *mPing* and *Ping16A* terminal sequence (1-40 bp). The SNP between *mPing* and *Ping16A* at position 16 (+16G/A SNP) is indicated by the red arrow. **b**, Transposition frequency of *mPing* variants that have mutations at the 5’ end in the yeast assay. X axis indicates *mPing* variants with mutations at 14 positions in the 5’ TIR and two positions outside the TIR. For example, *mPingG16A* represents an *mPing* variant having a G-to-A mutation at position 16. A variant *mPingC4A* was not included because the lack of qualified experiments. Y axis shows transposition frequency that was measured as *mPing* excision events per million cells and normalized to the control *mPing*. The error bars show standard deviation of 2-9 independent biological replicates. Letters above the bars indicate significant differences of transposition frequency between *mPing* variants and control (adjusted *P* value ≤ 0.05). The adjusted *P* values are based on a one-way ANOVA (*P* value = 2.37e-15, *F* value = 12.34, DF = 16) followed by a Tukey’s honest significant difference (Tukey’s HSD) test.

Prior studies identified four types of *mPing* elements (*mPingA-D*) in domesticated rice (Supplementary Fig. 4)^1^, representing four distinct deletion derivatives of *Ping*. Two of the four types (*mPingA,B*) were previously detected in *O. rufipogon* strains^14,15^. Here we detected all four types of *mPing* elements in *O. rufipogon* strains (Supplementary Table 4) indicating that *mPingA-D* arose prior to domestication in *O. rufipogon*. Like *mPing*, none of the 3,000 genomes analyzed in this study have more *Ping* elements (7-10) than the four strains undergoing *mPing* bursts^4^. *Ping* elements were detected in only 199 of 3,000 strains (6.6%) (Fig. 2 and Table 1) with most of the 199 (74.8%) having only a single copy and two strains having 4 *Pings* (Fig. 2b). In contrast, *Ping* elements were detected in 21 of 48 (43.7%) of the *O. rufipogon* strains analyzed (Table 1 and Supplementary Fig. 2). These data suggest that it is likely that *Ping* was selected against or lost from most strains during the hypothesized two or more domestication events from *O. rufipogon* populations^6,16^.

**Figure 4.**
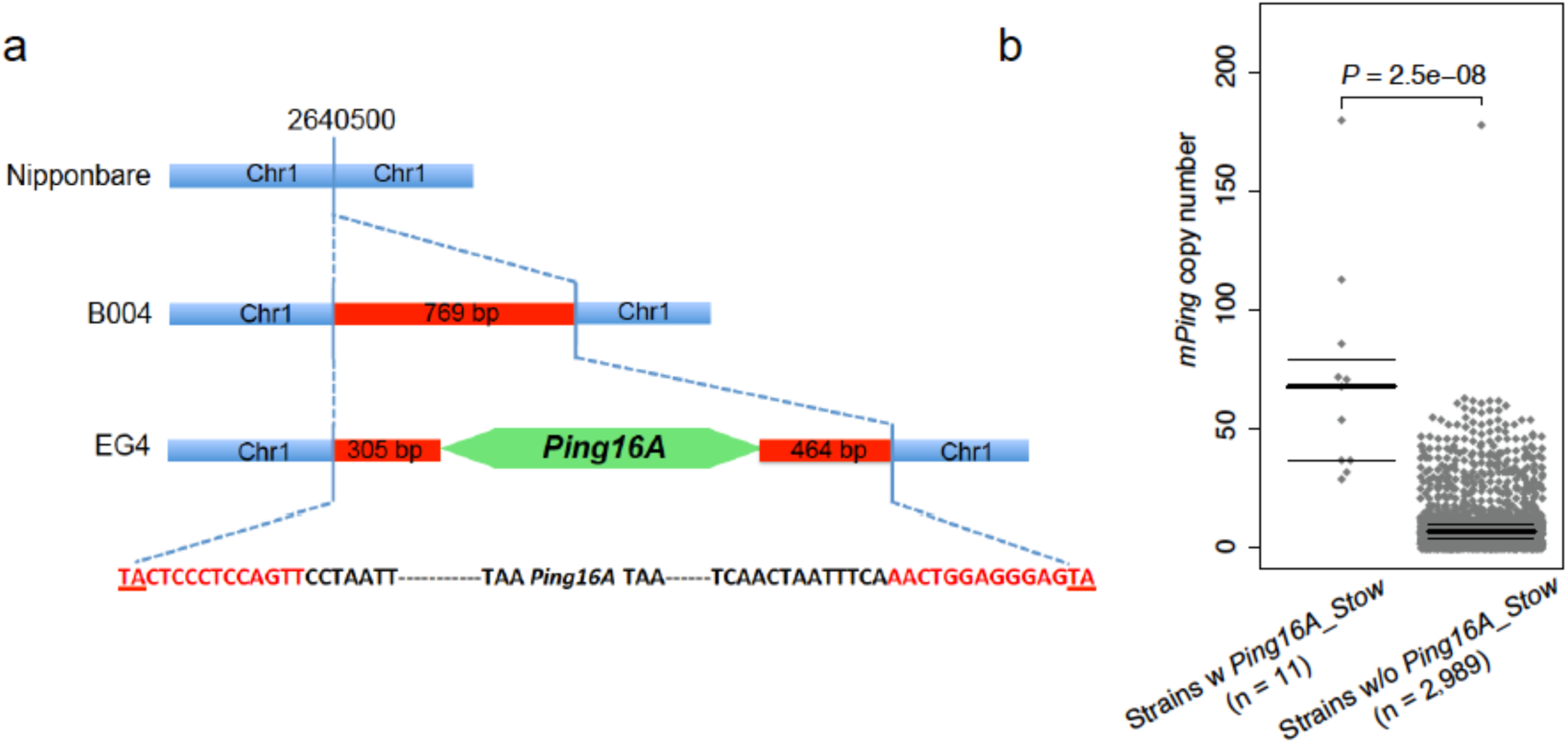
A *Ping* locus is associated with increased *mPing* copy number. **a,** Structure of the *Ping16A_Stow* insertion site. The *Ping16A* element (green arrow) is inserted in the middle of a non-autonomous *Stowaway* element (red box), which is not in Nipponbare (blue bar). The nucleotides shown within the blue dotted line are the sequences of the nonautonomous *Stowaway* element. **b**, Comparisons of *mPing* copy number in 3,000 rice strains with or without *Ping16A_Stow* in the genome. Gray dots indicate *mPing* copy number of rice strains in each category. Median and first/third quartiles of *mPing* copy number in each category are indicated by thick and thin black bars, respectively. The differences of *mPing* copy number between two categories are tested by a two-tailed Wilcoxon-Mann-Whitney test.

### Origin of a *Ping* variant and its possible significance

Analysis of the extensive collection of rice genomes revealed that a SNP distinguishing *Ping* and *mPing* (+16G/A) located adjacent to the 15-bp terminal inverted repeat (TIR) (Fig. 3a) and may be implicated in *mPing* bursts. *Pings* having these SNPs are referred herein as *Ping16G* (identical shared sequences with *mPing*) and *Ping16A*. First, all 21 *O. rufipogon* strains with *Ping* have only *Ping16G* which has the same sequence at +16G/A as *mPing* (Table 1). Thus, *Ping16G* is the original *Ping* and all 4 *mPing* types (*mPingA-D*, Supplementary Table 4) arose prior to domestication by internal deletion. Second, of the 199 domesticated rice strains with *Ping*, 31 have *Ping16G* while 154 have *Ping16A* (Table 1). The presence of the derived *Ping16A* in both *indica* and *japonica* strains was initially confusing as it suggested the unlikely scenario that this variant arose independently during the hypothesized two domestication events that led to these subspecies^6,16^. However, closer examination revealed that, where a determination could be made, all of the *Ping16A* loci in *indica* and admixed strains originated by introgression from *japonica* (Table 1). Thus, *Ping16A* has experienced limited but significant proliferation during and after *japonica* domestication such that it now accounts for the majority of *Ping* elements present in domesticated rice strains (Table 1).

### Reduced mobility of *Ping16A* in yeast assays

The TIRs and adjacent sequences of several DNA transposons have been shown to be functionally significant with mutations of these sequences reducing transposition frequency by decreasing the binding of transposase^17,18^. Because the SNP distinguishing *Ping16A* from *Ping16G* is adjacent to the 15-bp 5’ TIR (Fig. 3a), we employed a yeast assay to assess transposition rates of fourteen mutations within and two mutations adjacent to the 5’ TIR (Fig. 3b). In this assay, *Pong* transposase and an enhanced *Ping*_ORF1 (the putative binding domain) catalyzes transposition of *mPing* inserted in an ADE2 reporter gene, thereby allowing growth of yeast cells^11,19^. The results indicate that both the mutations adjacent to the TIRs (G16A and G17T) and 12 of 14 mutations in the TIR significantly reduced *mPing* transposition (Fig. 3b), supporting the hypothesis that this SNP (+16G/A) may have functional significance by reducing *Ping16A*’s mobility. Although *Pong* transposase was used in this experiment to facilitate the yeast transposition assays, its catalytic activity is almost indistinguishable from *Ping* transposase^19^. Furthermore, the reduced transposition of the G16A mutant (*mPingG16A*) was independently confirmed using *Ping* transposase (Supplementary Fig. 5).

### A *Ping* locus correlates with higher *mPing* copy number

The four strains previously shown to be undergoing *mPing* bursts (HEG4, EG4, A119, A123) have many (7-10) *Pings*, and all share only a single *Ping, Ping16A_Stow*^4^. This correlation suggests that acquisition of *Ping16A_Stow* may have initiated the burst. *Ping16A_Stow*, located on chromosome 1 (2640500-2640502), is comprised of the *Ping16A* variant inserted in a 769-bp *Stowaway* element (Fig. 4a). Of interest was whether any of the 3,000 strains had *Ping16A_Stow* and, if so, did they also have more *mPings*.

Among the 3,000 strains, 11 have *Ping16A_Stow* (188 have only the *Stowaway* insertion at this locus) (Table 1) and these strains have significantly more *mPings* (Two-tailed Wilcoxon-Mann Whitney test, *P* = 2.5e-08; Fig. 4b, Table 2, and Supplementary Table 5), providing additional correlative evidence for the involvement of *Ping16A_Stow* in *mPing* bursts.

### *Pong* has been stably silenced since domestication

*Pong* encoded proteins catalyze transposition of *mPing* in yeast and *A. thaliana* assays^10,11^ and in rice cell culture^1^. However, because *Pong* elements are epigenetically silenced in Nipponbare and in strains undergoing *mPing* bursts (HEG4, EG4, A119, A123)^4^, there is no evidence to date that *Pong* has an impact on *Ping* or *mPing* copy number or distribution.

Data from this study extend previous findings and suggest that *Pong* was silenced in *O. rufipogon* and has been stably silenced in domesticated rice. *Pong* elements are present in the genomes of almost all of the analyzed rice strains (99.1%, 2,972/3,000), and *Pong* copy numbers vary little within or between subgroups (Supplementary Fig. 6). On average, rice strains have four *Pong* elements (Fig. 1d). All *O. rufipogon* strains have *Pong* elements (Supplementary Fig. 2), except four (W1849, W1850, W2022, W2024), which appear to contain only *Pong* deletion derivatives (see Methods). As in domesticated rice, there is minimal *Pong* copy number variation among the *O. rufipogon* strains examined (Supplementary Fig. 2).

Six rice strains with higher *Pong* copy numbers (14-25) were analyzed to determine if this resulted from *Pong* activation. First, because active *Pong* elements produce proteins that catalyze *mPing* transposition, we tested if the genomes of these lines contained more *mPings*. However, all six strains had the same range of *mPing* copies as strains with few *Pongs* (Supplementary Table 6). Second, because host regulatory mechanisms suppress transposition, other potentially active TEs (elements shown previously to transpose when epigenetic regulation is impaired) may have been activated in these strains along with *Pong*. However, the six strains harbored average copy numbers of nine potentially active TEs (Supplementary Table 6). Taken together these data suggest that these six strains have accumulated silenced *Pong* elements during domestication. Finally, additional evidence for the stability of *Pong* silencing can be inferred from the observation that none of the 2,801 strains lacking *Ping* have a higher *mPing* copy number than strains with *Ping*.

### Discussion

Results of the evolutionary inventory of the members of the *mPing/Ping/Pong* TE family in wild and domesticated rice genomes suggest the following scenario for the origin of the *mPing* burst. All *mPing* subtypes in domesticated strains (*mPingA-D*) were generated prior to domestication, probably in *O. rufipogon,* by internal deletion from *Ping16G*. Furthermore, *Ping16G*, but not *Ping16A*, was detected in 21 of 48 *O. rufipogon* strains. The fact that only 31 of the 3,000 extant domesticated strains examined have *Ping16G* suggests that there has been a massive loss of this element during domestication. In contrast, the *Ping16A* variant was identified in the majority of the domesticated strains with *Ping* (154/199 strains). Its absence in *O. rufipogon* genomes indicate that it was either very rare in wild populations or that it arose during *japonica* domestication. During *japonica* domestication *Ping16A* has experienced limited but significant proliferation and has even been introgressed into a small number of *indica* strains (Table 1). Taken together these data indicate that *Ping16A* has become more widely distributed in domesticated strains, while *Ping16G* is disappearing.

Yeast assays testing the functional impact of several mutations in and adjacent to the *Ping* TIR demonstrate that the +16G (*Ping16G*) to +16A (*Ping16A*) polymorphism significantly reduces transposition frequency. Thus, *Ping16A* encoded proteins (which are identical to *Ping16G* encoded proteins) are more likely to catalyze the transposition of *mPing* (with its +16G) than of *Ping16A*. This situation is reminiscent of other autonomous elements that harbor sequences that reduce transposition frequency^20,21^. It has been hypothesized that autonomous TEs enhance their survival by evolving self-regulating mutations that reduce both host impact and epigenetic detection and silencing^21^.

The vast majority of strains with *Ping16A* have only one *Ping* (105/154 strains) and a moderate number of *mPing* elements (mean = 28). One of these strains is the reference strain Nipponbare where the inability to detect transposition of *Ping* or *mPing* was initially attributed to *Ping* silencing^22^. In fact, *Ping* is not silenced in Nipponbare nor in any other strain analyzed to date^4^. Rather it is transcribed and catalyzes (infrequent) transposition of *mPing*^2,4^. We speculate that strains with a single copy of *Ping16A* may be experiencing a balance, perhaps under stabilizing selection, between host survival and the maintenance of an active TE family in the genome.

The hypothesized balance between *Ping16A* and *mPing* elements and the host was perturbed in the subset of temperate *japonica* strains experiencing *mPing* bursts^4^ and it was suggested that the shared *Ping16A_Stow* locus may have been responsible^4^. Based on the evolutionary inventory presented in this study, it follows that *Ping16A_Stow* was generated in a temperate *japonica* strain when *Ping16A* transposed into a *Stowaway* element on chromosome 1. The *Stowaway* element was also present at this locus in *O. rufipogon* (Table 1). It is unlikely that this *Stowaway* is active as there are only 4 family members, each with less than 96% sequence identity, in the Nipponbare genome. Here we find that *Ping16A_Stow* is also shared by 5 of the 6 strains with the highest *mPing* copy numbers among the 3,000 strains analyzed (Table 2). The sixth strain, IRIS_313_15904, has a region of introgessed *indica* alleles at this location, which may have replaced the *Ping16A_Stow* locus in prior generations. The association of *Ping16A_Stow* with higher *mPing* copy numbers is consistent with its suggested role in triggering *mPing* bursts. However, the mechanism by which *Ping16A_Stow* may initiate the burst is unknown and warrants further investigation. Prior studies indicated that increased *Ping* transcripts were correlated with more *mPing* transpositions in strains undergoing *mPing* bursts^4,22^. Our unpublished data suggests that *Ping16A_Stow* does not produce more transcripts compared to other *Ping* elements, suggesting that mechanisms other than an increased transcript level from this locus may be responsible.

In conclusion, our data suggests that the key events of the burst, increased distribution of *Ping16A* and creation of the *Ping16A_Stow* locus, occurred during domestication. Other studies have shown that domestication can be associated with the loss of epigenetic regulation^23^, which may lead to the activation of TEs. However, our data indicate that *Pong* element copy number has been stably maintained from the wild ancestor through the generation of the thousands of domesticated strains, suggesting that epigenetic regulation was unaffected. In contrast, *Ping* activity has been sustained during domestication, resulting in the spread and amplification of the *Ping16A* variant and the generation of the *Ping16A_Stow* locus. Yet, the spread of *Ping* activity associated with exceptional *mPing* activity has been very limited in rice, likely due to the high level of self-fertilization, a domestication syndrome that has been observed in many flowering plants^24^.

## Materials and Methods

### Dataset

Illumina DNA sequencing reads of 3,000 rice strains were obtained from NCBI SRA project PRJEB6180. The metadata incorporating name and origin of the 3,000 rice strains was extracted from previously published Tables S1A and S1B^13^. The raw reads of 48 *O. rufipogon* strains were obtained from NCBI SRA under project accession numbers listed in Supplementary Table 2. The metadata associated with the subgroup classification of these 48 *O. rufipogon* strains was extracted from prior studies^6,26^. The raw reads of wild rice *Oryza glaberrima, Oryza glumipatula*, and *Oryza meridionalis* were obtained from NCBI SRA projects accession numbers SRR1712585, SRR1712910, and SRR1712972.

### Population structure and ancestral component analysis

The genotyped SNP dataset (release 1.0 3K RG 4.8 million filtered SNP Dataset) of the 3,000 rice genomes was obtained from SNP-Seek Database^27^ (http://snp-seek.irri.org). A subset of 270,329 SNPs was selected by removing SNPs in approximate linkage equilibrium using plink v1.09 (--indep-pairwise 1000kb 20kb 0.8)^28^. Population clustering analysis was performed by ADMIXTURE v1.3.0^29^ with K from 2 to 10. Most rice strains clustered into five subgroups (*indica*: IND, *aus*/*boro*: AUS, *aromatic* (*basmati*/*sadri*): ARO, temperate *japonica*: TEJ, and tropical *japonica*: TRJ) when K is 5. Using the ancestral analysis of ADMIXTURE under the K = 5 model, a rice strain was assigned to one of these five subgroups if it had more than 80% of its ancestral component from a given subgroup. Any strains that had no major ancestral component (< 80%) were categorized as admixed (ADM) strains. During the preparation of this study Wang et al. published an analysis of the same dataset^16^. The subgroup classifications were compared between the two studies and the results are consistent except that Wang et al. identified additional subgroups in *indica* and *japonica*.

The 4.8 million filtered SNPs were imputed and phased with BEAGLE v5.0^30^ using default parameters. A total of 768 strains with major ancestral component over 99.99% were used as reference panels for five rice subgroups (344 *indica* strains, 111 *aus/boro* strains, 31 *aromatic* strains, 124 temperate *japonica* strains, and 158 tropical *japonica* strains). Local ancestry assignment was performed on strains of interest with RFMix v2.03^31^ using default parameters.

### Copy numbers characterization

The *mPing, Ping,* and *Pong* insertion sites across the 3,000 rice genomes were genotyped using RelocaTE2 (aligner = BLAT mismatch = 1 len_cut_trim = 10)^12^. Element-specific sequence differences were identified and used to distinguish *Ping* and *Pong* from *mPing* insertions (Fig. 1a). Three separate runs of RelocaTE2 were performed using *mPing*, *Ping*, and *Pong* as queries. Paired-end reads where one read of a pair matched the internal sequence of a *Ping* element (253-5,164 bp) and the mate matched to a unique genomic region of the Nipponbare reference genome (MSU7) were used to differentiate *Ping* insertions. Similarly, paired-end reads where one read matched the internal *Pong* element sequence (23-5,320 bp) and the mate matched to a unique genomic region of MSU7 were used to identify *Pong* insertions. An equivalent approach was undertaken with *mPing* sequences but the prior identified *Ping* and *Pong* insertion sites were removed from the *mPing* RelocaTE2 results to generate final *mPing* insertions. RelocaTE2 analysis was performed in 48 *O. rufipogon* genomes to identify *mPing, Ping*, and *Pong* insertions. However, the short insert size and insufficient read depth of *O. rufipogon* sequencing libraries prevented distinguishing *Ping* and *Pong* insertions from *mPing*.

Copy numbers of *mPing, Ping,* and *Pong* elements were estimated from the ratio of the element sequence coverage to the genome-wide average sequence coverage. All sequencing reads associated with a given repeat element were extracted from the RelocaTE2 results. The reads were aligned to the element using BWA v0.7.12^32^ with default parameters. Alignments with less than 2 mismatches were retained for further analysis. The sequence coverage of each position in the element was calculated using mpileup command in SAMtools v0.1.19^33^. The average sequence coverage of *mPing* in each genome was calculated from the average read depth of positions 1-430, while *Ping* and *Pong* coverage were calculated using the average read depth of positions 260-3,260 so that unique regions in the targeted element were considered for the assessment. The genome-average sequence coverage of each genome was calculated using qualimap v2.1.2^34^.

### Analysis of *Ping16A_Stow*

The pre-aligned BAM files of 3,000 rice genomes (http://s3.amazonaws.com/3kricegenome/Nipponbare/”Strain_Name”.realigned.bam) were analyzed to determine if a *Stowaway* element was present at the *Ping16A_Stow* locus Chr1:2640500-2640502. A total of 199 rice genomes with signatures of TE insertions at the *Ping16A_Stow* locus (reads with only partial “soft clipped” alignments) were analyzed to confirm the *Stowaway* element insertion. A pseudogenome was built of a single *Stowaway* element and its 2 kb flanking sequences of the position Chr1:2640500-2640502. The sequencing reads from each of the 199 rice genomes were aligned to the pseudogenome using BWA and SAMtools with default parameters followed by analysis of the BAM files to identify junction reads covering both the *Stowaway* and its flanking sequence. All of these 199 strains were confirmed to have the *Stowaway* element at the position Chr1:2640500-2640502.

A similar approach that identifies the *Stowaway* element insertion was used to identify *Ping* insertions in the *Stowaway* element at the *Ping16A_Stow* locus. A pseudogenome was built using a *Ping* element and its flanking sequences, which are 1-305 bp of the *Stowaway* element upstream *Ping* insertion and 306-770 bp of the *Stowaway* element downstream *Ping*. The sequencing reads of these 199 rice genomes were aligned to the pseudogenome using BWA and SAMtools with default parameters. Analysis of junction reads that cover both *Ping* element and its flanking *Stowaway* element identifies eleven strains having a *Ping* insertion in the *Stowaway* element at the *Ping16A_Stow* locus (Supplementary Table 5).

### Analysis of +16G/A SNP genotype

A locus-specific approach was used to analyze the genotype of the +16G/A SNP on the *Ping* element in rice. *Ping*-containing reads of each locus were extracted from the RelocaTE2 results. The reads were aligned to the Nipponbare *Ping* element using BWA with default parameters. Alignments with less than 2 mismatches were analyzed using mpileup command in SAMtools to generate a read depth profile, which includes base composition information at each position. The nucleotide counts at the +16G/A SNP were obtained from the read depth profile. A *Ping* with more than two reads supporting G was genotyped as *Ping16G*, while a *Ping* locus with more than two reads supporting A was genotyped as *Ping16A*.

For *O. rufipogon,* all reads aligning to *mPing, Ping*, and *Pong* were pooled to analyze the base composition at the +16G/A SNP because *mPing, Ping*, and *Pong* insertions could not be efficiently sorted. An *O. rufipogon* genome was categorized as a genome having *Ping16G* or *Ping16A* based on whether they had more than two reads supporting G or A. Strains that have more than two reads supporting both G and A were further analyzed to clarify whether the *Ping16A* is present in these genomes. For example, strain W1230 had both G (288 reads) and A (23 reads) at +16G/A SNP. These A-supporting reads and their mates were extracted from W1230 sequences and aligned to pseudogenomes that have W1230 *mPing* or *Ping* inserted in MSU7. All of these A-supporting reads were uniquely aligned to an *mPing* locus Chr3:25526483-25526485. This *mPing* locus contains a 430 bp *mPingC* element that was successfully assembled from locus-specific paired-end reads, suggesting these A-supporting reads were from *mPing* not *Ping*.

### Assembly and classification of *mPing* sequences

A locus-specific assembly was performed to recover full-length *mPing* sequences from rice sequences. The sequencing reads matching *mPing* were obtained using RelocaTE2, assembled using velvet v1.2.09 (MAXKMERLENGTH = 31 -ins_length 400-exp_cov 50 -scaffolding yes)^35^. The flanking non-*mPing* sequences were removed from the assembled sequences. Any *mPing* candidate loci containing sequence gaps were removed from the analysis. The remaining full-length *mPing* sequences were compared using BLAST v2.2.26 to build an undirected graph with python package NetworkX (https://networkx.github.io). Each node in the graph is an *mPing* sequence and each edge is a connection, which requires two *mPing* sequences are properly aligned (number of gaps or mismatches ≤ 4). The *mPing* sequences in each subgraph represent a class of *mPing*. Representative sequences were extracted from each *mPing* class and aligned with four canonical defined *mPing* classes (*mPingA, mPingB, mPingC*, and *mPingD*) from the prior study^1^ using MUSCLE v3.8.425^36^ with default parameters. The multiple sequence alignment in MSA format was converted into VCF format using msa2vcf.jar tool (https://github.com/lindenb/jvarkit) to identify polymorphic sites. The assembled *mPing* sequences were classified into classes based on their breakpoints and point mutations compared to the four canonical *mPing* classes.

The reads of *O. rufipogon* strains were aligned to four canonical defined *mPing* classes (*mPingA, mPingB, mPingC*, and *mPingD*) using BWA with default parameters. Alignments with less than 2 mismatches were manually inspected using Integrative Genomics Viewer (IGV) v2.3.0^37^ to determine if the reads cover breakpoint of each *mPing* class in each strain. A strain with two or more reads covering the breakpoint of an *mPing* class was identified as a strain containing this *mPing* class.

### Phylogenetic analysis

The 270,329 SNPs used for ADMIXTURE analysis were used to genotype HEG4, EG4, A119, and A123 using GATK UnifiedGenotyper v3.4-46^38^. The phylogenetic tree of rice strains was built using a Neighbor-Joining method implemented in FastTree v2.1.10 (- noml -nome)^39^. The sequencing reads for the 48 *O. rufipogon* strains were analyzed to obtain a SNP dataset. Briefly, paired-end reads were aligned to MSU7 using SpeedSeq v 0.1.0^40^, which uses BWA to align reads, Sambamba^41^ to sort alignments, and SAMBLASTER^42^ to mark PCR duplicates. The resulting BAM files were analyzed with GATK UnifiedGenotyper to perform SNP calling. Filtering parameters were used to retain only homozygous SNPs that did not overlap repetitive sequences. These high-quality SNPs were extracted and converted into PHYLIP format multiple sequence alignment for phylogenetic analysis with RAXML v8.2.8^43^ under a GTRGAMMA model (- m GTRGAMMA). Bootstrap was performed using 100 iterations (-f a -# 100). Wild rice *O. glaberrima, O. glumipatula*, and *O. meridionalis* were treated as outgroups. Graphical representations of the phylogenetic trees were generated in R using “APE” libraries^44^.

### Yeast transposition assay

*mPing* was amplified with Phusion High-Fidelity PCR Master Mix (Thermo Fisher Scientific) using the control *mPing* primers (*mPing* F and *mPing* R) or mutation containing primers (i.e. *mPing* F and *mPing16A* R; Supplementary Table 7). The primary PCR products were then amplified with ADE2 TSD F and ADE2 TSD R primers (Supplementary Table 7) to add ADE2 homologous sequences. Purified PCR products were co-transformed into *Saccharomyces cerevisiae* strain JIM17^45^ with *Hpa*I digested pWL89a plasmid as described in the prior work^46^. Plasmids were isolated from yeast strains using the Zymo Yeast Plasmid Miniprep kit (Zymo Research) and transformed into *Escherichia coli* for sequence validation.

Sequence verified plasmids were transformed into *S. cerevisiae* strain CB101^45^ containing previously described pAG413 GAL ORF1 Shuffle1 NLS and pAG415 GAL *Pong* TPase L384A, L386A plasmids^19^. The transposition rate was measured as described in the prior study^11^. Briefly, 3 ml cultures were grown in CSM-His-Leu-Ura (dextrose) for 24 h at 30°C, and 100 µl was plated onto 100 mm CSM-His-Leu-Ura-Ade (galactose) plates. The total number of yeast cells was calculated by plating a 10^−4^ dilution of the cultures onto YPD plates. The numbers of colonies on the galactose plates were determined after 10 days of incubation at 30°C. The transposition rate was determined by dividing the galactose colony count by the total number of cells plated.

### Statistical analysis

Sample sizes, statistical tests, and *P* values are indicated in figures or figure legends. Linear regression, two-tailed Pearson’s correlation, two-tailed Wilcoxon-Mann-Whitney, one-way ANOVA and Tukey’s honest significant difference (Tukey’s HSD) test were performed with lm, cor.test, wilcox.test, aov, and TukeyHSD functions in R.

### Code availability

RelocaTE2 and other code used in this study are available at https://github.com/stajichlab/Dynamic_rice_publications or https://doi.org/10.5281/zenodo.1344714.

## Acknowledgments

We thank Drs. Hongru Wang and Shujun Ou for discussions of data analysis and Dr. Shaohua Fan and Julia Adams for valuable comments on the manuscript. This work was supported by National Science Foundation grants (IOS-1027542 to S.R.W. and J.E.S. and MCB-1651666 to C.N.H.). Data analyses were performed on the UC Riverside High-Performance Computational Cluster supported by National Science Foundation grant DBI-1429826 and National Institutes of Health grant S10-OD016290.

## Author contributions

J.C., J.E.S., and S.R.W. conceived the study. J.C. and L.L. analyzed the sequence data. J.B., S.D., and C.N.H. performed the yeast experiment and analyzed the data. J.C., C.N.H., J.E.S., and S.R.W. wrote the paper.

## Competing interests

The authors declare no competing financial interests

